# Structural Basis for the Activation of Proteinase-Activated Receptors (PARs) by Endogenous Ligands

**DOI:** 10.1101/2024.07.08.602624

**Authors:** Zongyang Lyu, Xiaoxuan Lyu, Guliang Xia, Daniel Carney, Vinicius M. Alves, Matthew Falk, Nidhi Arora, Hua Zou, Aaron McGrath, Yanyong Kang

## Abstract

The proteinase-activated receptor (PAR) subfamily of G protein-coupled receptors (GPCRs) include four members, PAR1-PAR4, that play critical roles in hemostasis, thrombosis, embryonic development, wound healing, inflammation, and cancer progression. The PARs share a unique activation mechanism driven by proteinase cleavage at a specific site within the extracellular amino-terminus, exposing a ‘tethered ligand’ that self-activates the receptor. Subsequent activation allows PAR family members to initiate complex intracellular signaling networks via traditional G protein-mediated pathways and beta-arrestin signaling and, in this way, the PARs link extracellular protease signaling molecules to cellular functions. Despite a primary reliance on biochemical studies for understanding tethered ligand recognition, direct structural visualization of these ligand-receptor complexes has been elusive. Here, we present structural snapshots of activated PAR1 and PAR2 bound to their endogenous tethered ligands, revealing, for the first time, shallow and constricted orthosteric binding pockets and highlighting critical residues involved in ligand recognition and receptor activation. Surprisingly, comparisons with antagonist-bound structures show minimal conformational changes in the TM6 helix, a typical signature of GPCR activation, with large movements of TM7 observed upon activation. These insights lead to the identification of a common mechanism for PAR1 and PAR2 activation and provide a structural template for designing novel antagonists targeting the orthosteric binding site, potentially opening new avenues for therapeutic interventions.

## Introduction

Proteinase-activated receptors (PARs) represent a unique subgroup within the broader family of G protein-coupled receptors (GPCRs), which are essential to myriad biological processes. These receptors are activated by proteolytic cleavage by various serine proteases, unveiling a “tethered ligand” within the receptor structure itself. This tethered ligand subsequently binds intramolecularly to initiate signaling, distinguishing PARs from other GPCRs that are typically activated intermolecularly by soluble ligands[1]. Notably, PAR1 and PAR2 are crucial for their roles in human physiology and pathology [2–4].

PAR1, the earliest discovered member of this family, has been extensively studied due to its vital role in hemostasis and thrombosis[5, 6]. Activated primarily by thrombin, a critical enzyme in the coagulation cascade[7], these activities underscore PAR1’s pivotal role in vascular biology and as a therapeutic target for novel antithrombotic therapies that aim to reduce thrombotic risk without the associated bleeding risks present in other anticoagulants[8].

Conversely, PAR2 is mainly activated by trypsin and tryptase, playing a varied role in inflammation and immune response[9]. Unlike PAR1, its functions are implicated in conditions such as inflammation, pulmonary disorders, and cancer progression[10, 11]. The ability of PAR2 to respond to multiple proteases suggests its broader physiological implications and complexity, making it an appealing target for pharmaceutical development aimed at various inflammatory and autoimmune disorders[12].

Advancing from this foundational knowledge, our research delves into the structural analysis of PAR1 and PAR2, revealing their activation mechanisms in unprecedented details. We have resolved the structures of these receptors in complex with their endogenous tethered ligands, providing clear illustrations of receptor activation dynamics. Our analysis reveals that both PAR1 and PAR2 feature shallow and narrow orthosteric binding pockets essential for ligand recognition and receptor interaction. This structural characterization not only deepens our understanding of their molecular functionality but also identifies critical residues that are crucial for receptor activation. Surprisingly, our comparative studies with antagonist-bound structures reveal minimal conformational shifts in the TM6 helix, suggesting a subtle yet effective activation mechanism distinct from the dramatic transformations observed in other GPCRs. These insights are vital for comprehending the specificity of receptor activation and signaling.

Leveraging these structural discoveries, we propose a unified model for PAR activation. Our detailed understanding of ligand-receptor interactions and conformational states forms a structural foundation for designing novel agonists and antagonists that specifically target the orthosteric binding sites of PAR1 and PAR2. This strategy promises highly selective therapeutic options, enhancing treatment efficacy while minimizing off-target effects, thereby opening new therapeutic possibilities for diseases with underlying PAR2 activation or overactivity.

## Results and discussion

### Structure determination and overall structures

To elucidate the agonist-binding and activation mechanisms of PAR1 and PAR2, we focused on their natural agonist, the tethered ligand. We employed consistent strategies in designing constructs for both receptors. Briefly, PAR1 (residues 42-425) and PAR2 (residues 37-397) were cloned into the pFastbac vector just after the signal peptide (Extended Data Fig. 1). This arrangement simulates protease cleavage, thereby exposing the first amino acid of the tethered ligand, which is crucial for receptor activation. To stabilize the PAR1 and PAR2 complexes with Gαq proteins, we implemented the NanoBiT tethering strategy by attaching the large BiT (LgBiT) to the C-terminus of each receptor, followed by a Flag tag for purification purposes[13].

Additionally, we utilized an engineered Gαq protein (GqiN), in which the N-terminus (residue 1– 32) of Gαq was replaced with the N-terminus (residue 1–28) of Gαi[14]. This modification significantly enhances the protein’s affinity for the scFv16 antibody, a tool that has been instrumental in elucidating numerous receptor/G-protein complexes. The Gβ subunit in both the PAR1 and PAR2 complexes features a C-terminal HiBiT, which pairs with the LgBiT at the receptor’s C-terminus, thereby securely anchoring the G protein complex.

For assembling the PAR1-Gαq and PAR2-Gαq complexes, receptor subunits, Gαq, Gβ, Gγ, and scFv16 were co-expressed in Hi5 insect cells. The complexes were then purified using Flag affinity resin. The structures of the PAR1-Gαq and PAR2-Gαq complexes were resolved via cryo-EM to resolutions of 2.74 Å and 3.1 Å (Extended Data Fig. 2), respectively. The clarity of the cryo-EM density maps enabled accurate modeling of most side chains, including those of the tethered ligands, the receptors, the Gαβγ heterotrimer, and scFv16 (Figure 1A-F). Both complexes exhibit a highly similar overall structure, adopting a canonical seven-transmembrane helix domain (TMD), with root mean square deviation (RMSD) values of 1.07 Å over 205 Cα atoms. The tethered ligands bind at the orthosteric pocket formed by transmembrane helices TM5, TM6, TM7, the N-terminal loop, and extracellular loop 2 (ECL2).

**Figure 1.**
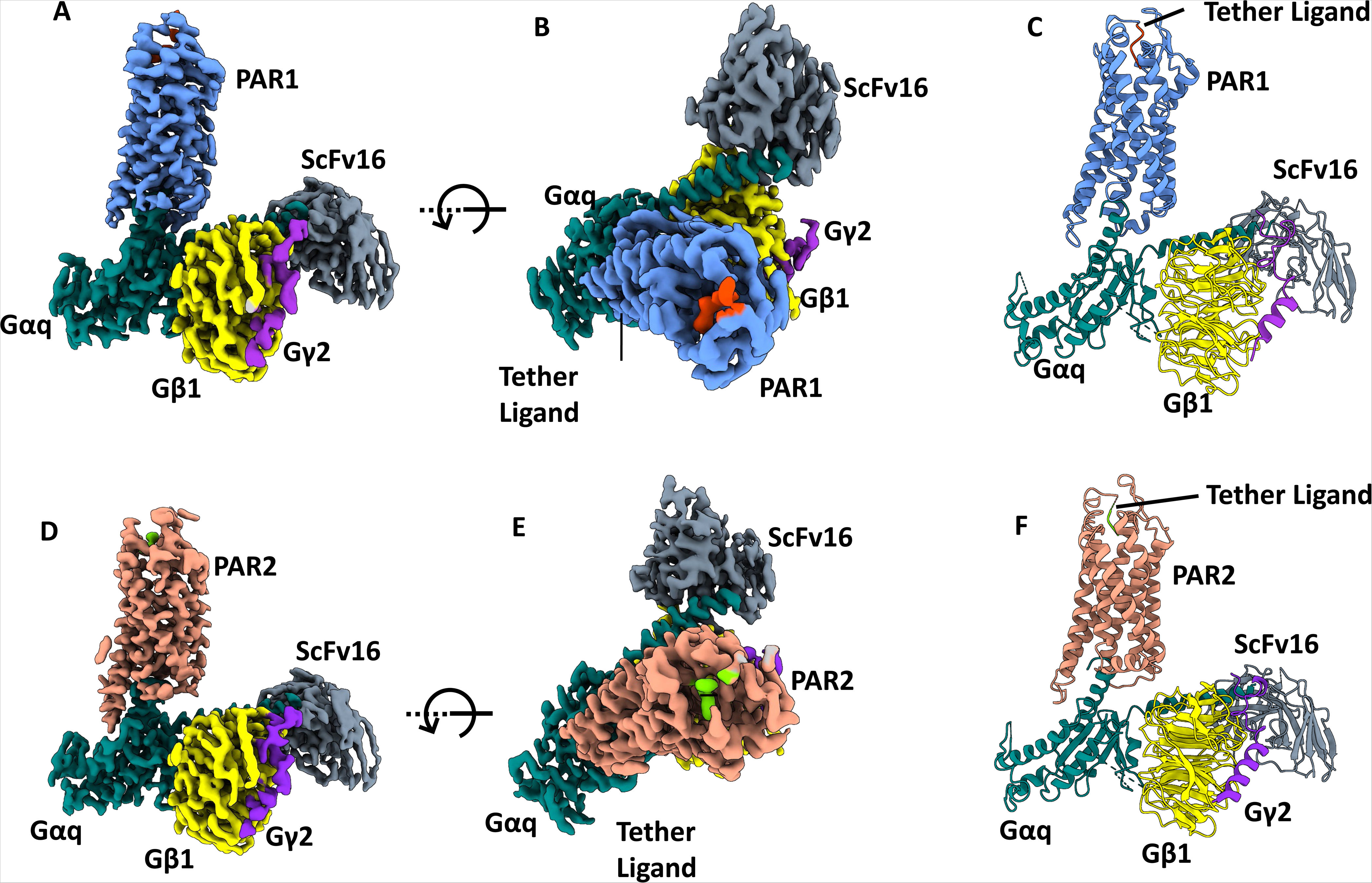
Cryo-EM structure of the PAR1-Gαq-scFv16 complex and PAR2-Gαq-scFv16 complex. **A,** Cryo-EM density map of PAR1-Gαq-scFv16 with tethered ligand. **B,** The same density map shown after rotating 90°. **C**, Model of PAR1-Gαq-scFv16 with tethered ligand. **D**, Cryo-EM density map of PAR1-Gαq-scFv16 with tethered ligand. **E**, The same density map shown after rotating 90°. **F**, Model of PAR1-Gαq-scFv16 with tethered ligand. (PAR1 tethered ligand, orange red; PAR1, blue; Gαq, teal; Gβ, yellow; Gγ, purple; scFv16, gray; PAR2 tethered ligand, lime; PAR2, coral)

### Tethered Ligand Binding and the Orthosteric Binding Pocket

PAR1 is uniquely activated by thrombin through a specialized proteolytic mechanism that sets it apart from most other G protein-coupled receptors. This activation begins when thrombin cleaves PAR1 at residue R41 located within its extracellular N-terminal domain[15]. This proteolytic event exposes a new sequence starting with SFLLR, which acts as a “tethered ligand.” Our PAR1-Gαq cryo-EM map reveals robust density for these initial five residues. They adopt an extended conformation that binds into the core of PAR1, making extensive contact with TM1, TM5, TM6, TM7, and the second intracellular loop (ICL2) (Figure 2A). Within this intricate interaction network, the tethered peptide is nestled into a shallow and narrow pocket, where the first residue, S42, forms extensive hydrogen bonds with residues H255 of ECL2, Y337 of TM6, and Y350 of TM7 (Figure 2B). Furthermore, F43 contributes critical van der Waals interactions with residues L96 of the N terminal loop, E347 of TM7, and Y350 of TM7, illustrating a complex network of interactions that facilitate the activation of the receptor (Figure 2C).

**Figure 2.**
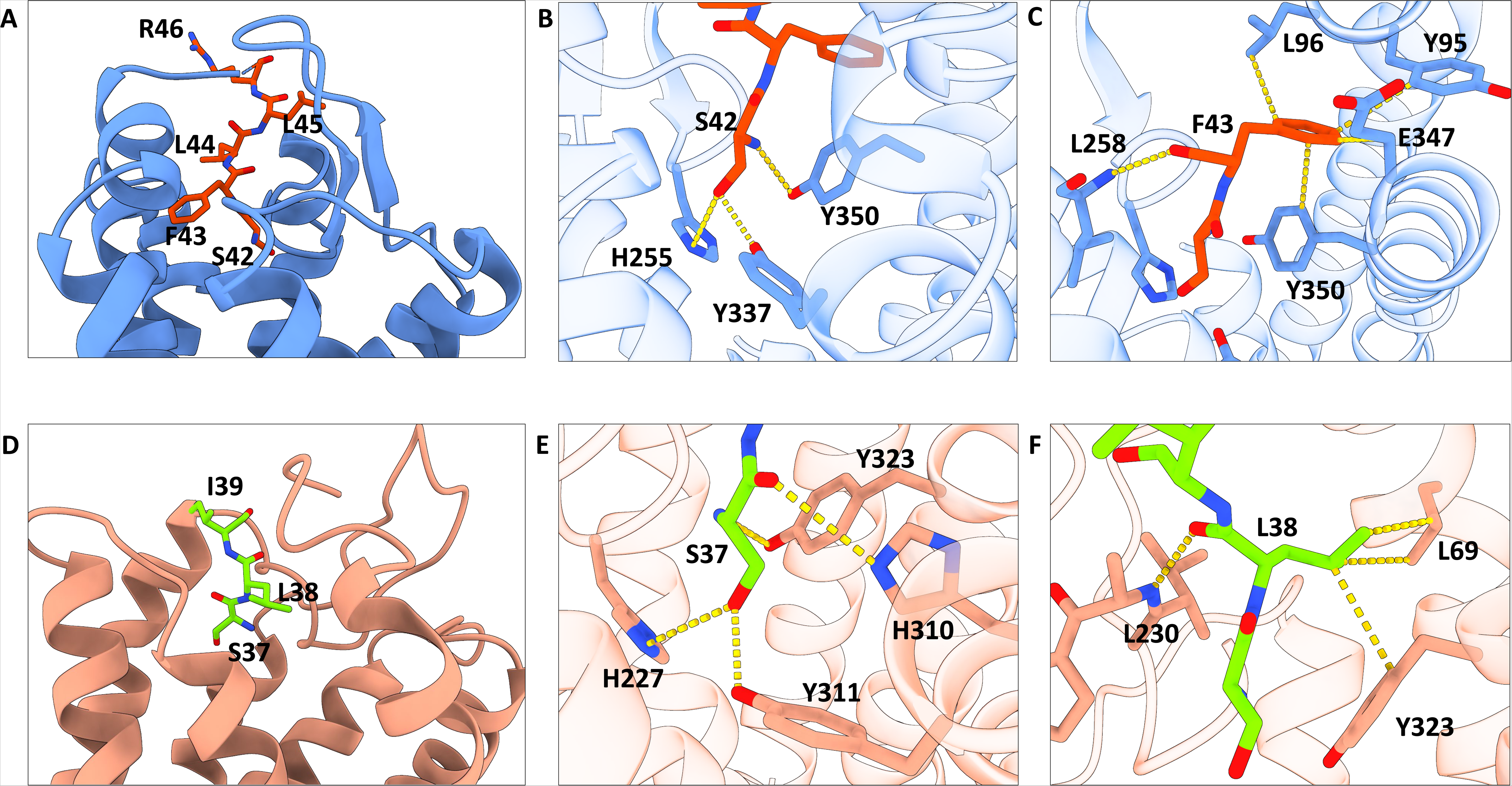
Tethered peptide ligand binding to the orthosteric pocket in PAR1 and PAR2. **A**, PAR1 tethered ligand SFLLR (orange) binds to PAR1 (blue); **B**, H-bond interactions between S42 of tethered ligand and PAR1; **C**, Details of interactions between F43 of tethered ligand and PAR1; D, PAR2 tethered ligand SLI (Lime) binds to PAR2 (Coral); **E**, H-bond interactions between S37 of tethered ligand and PAR2; **F**, Details of interactions between S37 of tethered ligand and PAR2.

In contrast, PAR2 activation involves a direct mechanism by trypsin, which cleaves the receptor at R36, thereby unveiling a new N-terminal sequence starting with SLIG that serves as a tethered ligand[16]. This ligand then folds back onto the receptor itself, binding intramolecularly and inducing significant conformational changes that are essential for receptor activation. Our cryo-EM mapping of the PAR2-Gαq complex has effectively resolved the initial three residues, SLI, showing a similar pattern of interaction as observed in PAR1. This tethered ligand also binds to a shallow pocket, engaging in key interactions with TM1, TM5, TM6, TM7, and ICL2 (Figure 2D). Mutations of these residues to alanine result in a significant reduction in agonist potency by more than 10-fold, highlighting the critical nature of these interactions for receptor functionality[17]. Notably, residues S37 and L38 play important roles in this binding configuration. S37 reaches into the narrow base of the pocket, having multiple hydrogen bonds with H227 of ECL2, Y311 of TM6, and H310 of TM6 (Figure 2E), while L38’s side chain is involved in a polar interaction with L230 of TM4 and forms van der Waals interactions with L69 of TM1 and Y323 of TM7, illustrating the detailed mechanism of ligand-receptor interactions that lead the activation of PAR2 (Figure 2F).

### Structural Comparison of Active and Inactive States of PAR1

The structure of PAR1 bound to the antagonist vorapaxar (PDB 3VW7) illustrates how vorapaxar occupies the orthosteric binding site of PAR1, effectively inhibiting its activation[18]. By forming hydrogen bonds and hydrophobic interactions with several residues within the binding pocket, vorapaxar stabilizes PAR1 in an inactive conformation and impedes the conformational changes necessary for G protein coupling. Comparative analysis with the inactive vorapaxar-bound PAR1 reveals key structural differences from the tethered ligand-bound active state of PAR1.

Firstly, vorapaxar functions as an antagonist by occupying the site where the tethered ligand’s first residue, S37, would typically bind. In the cryo-EM structure, S42 is positioned similarly to the lactone ring of vorapaxar, forming essential hydrogen bonds with nearby residues, which underscores its role in promoting agonist activity (Figure 3A-B, Extended Data Fig. 5). Secondly, unlike other Class A and B GPCRs, which typically exhibit significant activation-associated movements in TM6 (such as a 9 Å outward shift in rhodopsin[19] and about 16 Å in GLP1R when engaged with G proteins[20]), PAR1 shows a more modest 4.7 Å outward movement of TM6 in the active form compared to the vorapaxar-bound state. Furthermore, TM5 does not undergo dramatic extensions but does exhibit a 2.6 Å outward movement in the ligand-bound state compared to the inactive state, highlighting the unique activation dynamics of PAR1 (Figure 3A).

**Figure 3.**
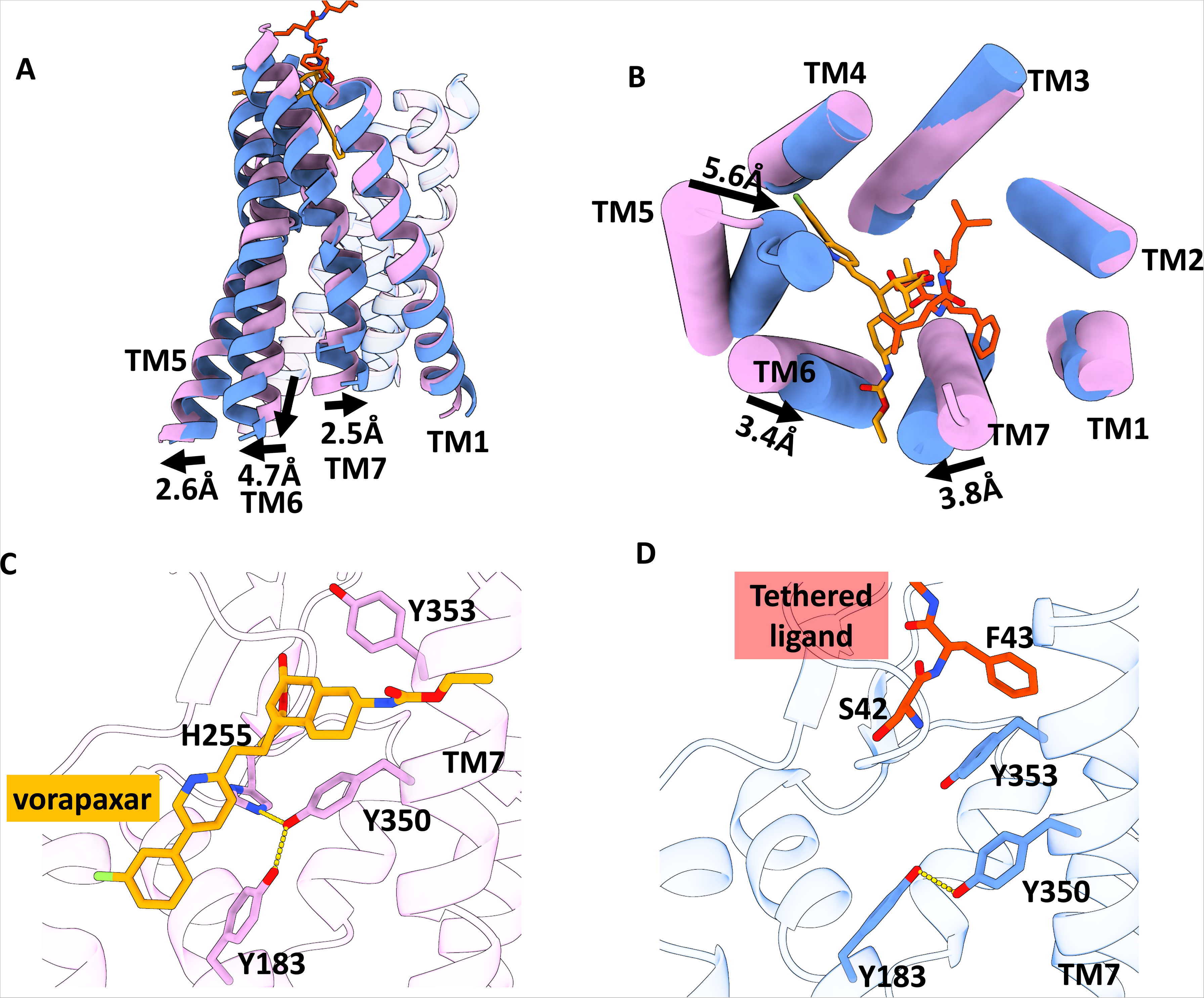
Structural Comparison of Active and Inactive States of PAR1. **A**, Side view comparison of the tethered ligand-bound to active PAR1 and the antagonist vorapaxar-bound to inactive PAR1 (PDB 3VW7). Active PAR1 is depicted in blue, and inactive PAR1 is shown in pink. **B**, Extracellular view of the comparison between tethered ligand-bound active PAR1 and vorapaxar-bound inactive PAR1 (PDB 3VW7). The tethered peptide is colored in orange, while vorapaxar is shown in yellow. **C**, Conformations of key residues in PAR1 bound to the antagonist vorapaxar. **D**, Conformations of key residues in PAR1 bound to the tethered ligand.

Thirdly, both vorapaxar and the tethered ligand bind close to the extracellular surface of PAR1, differing from other GPCRs where ligands typically penetrate deeper into the transmembrane core. In contrast with the vorapaxar-bound state, movements of TM6 and TM7 in the active form exceed 3Å towards each other, closing the tunnel that accommodates vorapaxar’s ethyl carbamate tail. The most notable change is an inward movement of TM5 by 5.6 Å, which narrows the pocket between TM4 and TM5 tailored for vorapaxar’s fluorophenyl pyridine tail (Figure 3B). The tethered ligand SLI penetrates perpendicularly into a small orthosteric pocket formed by TM1, TM5, TM6, TM7, and ICL2.

Lastly, significant conformational shifts induced by the tethered peptide are observed in Y350 and Y353. In the vorapaxar-bound structure, Y350 forms a strong hydrogen bond with vorapaxar, maintaining Y350 and Y353 of TM7 in a raised position (Figure 3C). However, the tethered ligand forces these residues downward. The sidechain of Y350 on TM7 engages in a polar hydrogen bond with Y183 of TM3, leading to the downward movement of TM6 and activating the receptor (Figure 3D).

### Structural Comparison of Active and Inactive States of PAR2

Previous work reveals distinct binding mechanisms of PAR2 antagonists. For instance, the antagonist AZ8838 (PDB 5NDD) binds to a fully occluded pocket near the extracellular surface, showing slow binding kinetics advantageous for competing against tethered ligands[21]. Another antagonist, AZ3451 (PDB 5NDZ), binds to a distant allosteric site outside the helical bundle, preventing necessary structural rearrangements for receptor activation and signaling[21, 22].

In our tethered ligand-bound PAR2 structure, the tethered ligand shows a very different binding site to other Class A or Class B GPCRs, close to the extracellular side of the receptor (Extended Data Fig. 4). Similar to the PAR1-Gαq complex, PAR2 exhibits minimal movement on TM6, with a 3.4 Å outward shift and around a 2 Å shift to the intracellular side. There is no extension on TM5, but it shows a 2.6 Å outward movement to accommodate the Helix-5 of the G protein (Figure 4A). Due to the different binding model between the tethered ligand and AZ8838, the tethered ligand induces a different conformation of TM6 around the binding pocket, causing a 2.1 Å shift (Figure 4B).

**Figure 4.**
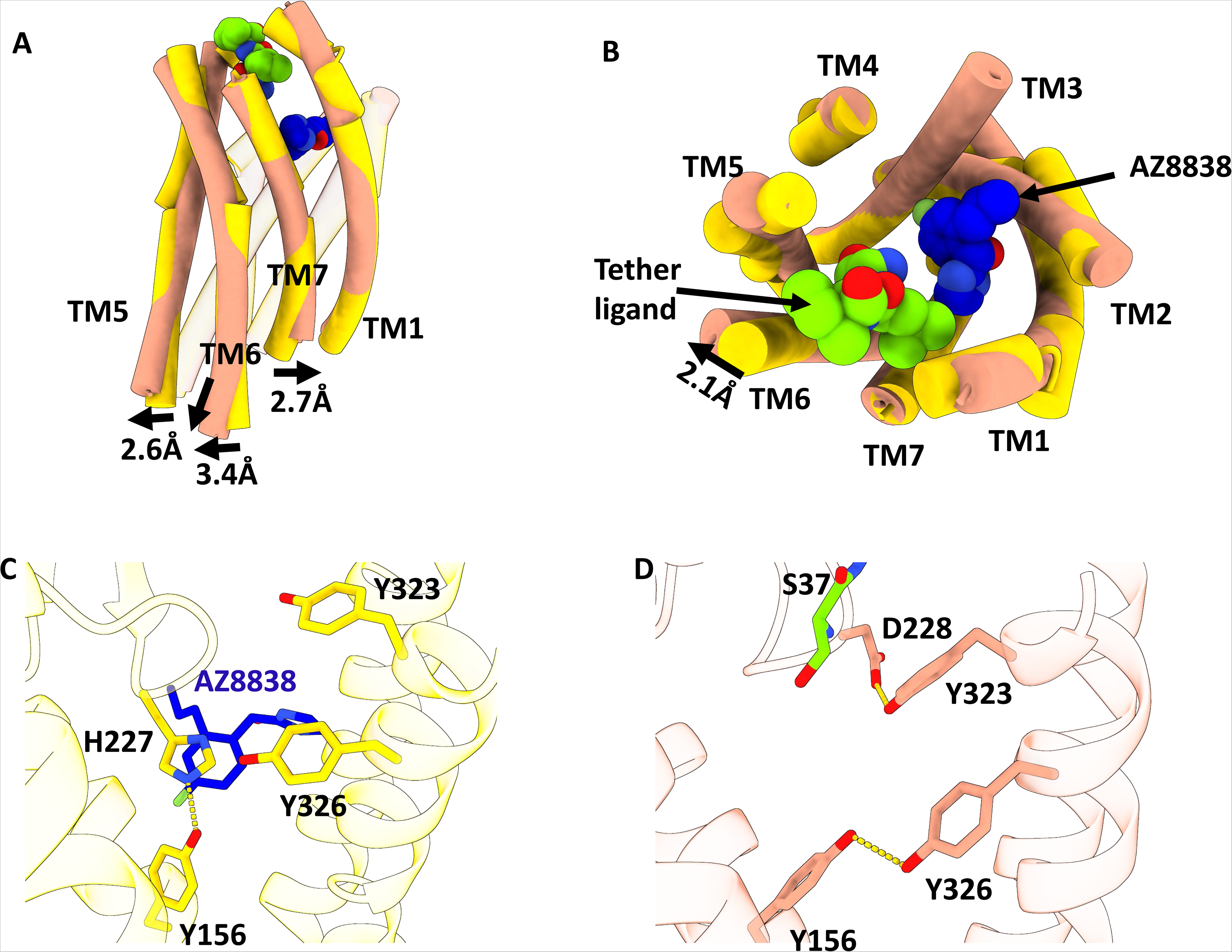
Structural Comparison of Active and Inactive States of PAR2. **A**, Side view comparison of the tethered ligand-bound active PAR2 and the antagonist AZ8838-bound inactive PAR2 (PDB 5NDD). Active PAR2 is depicted in coral, and inactive PAR1 is shown in yellow. **B**, Extracellular view of the comparison between tethered ligand-bound active PAR2 and AZ8838-bound inactive PAR2 (PDB 5NDD). The tethered peptide is colored in lime, while AZ8838 is shown in blue. **C**, Conformations of key residues in PAR2 bound to the antagonist AZ8838. **D**, Conformations of key residues in PAR2 bound to the tethered ligand.

The tethered ligand exhibits a different binding model from AZ8838, which is completely solvent inaccessible in a small, deep binding pocket below the orthosteric site (Figure 4C). Upon binding at the orthosteric pocket, the sidechain of Leu38 sterically pushes Y323 of TM7 downwards, forming a stabilizing H-bond with D228 of ECL2 via the sidechain hydroxyl group. This movement causes a significant downward shift of Y326 of TM7. Both residues are critical for activation, as shown in mutagenesis experiments (reference needed). The movement of Y326 of TM7 results in Y156 of TM3 moving away from H227 of ECL2, breaking an H-bond and forming a new stabilizing H-bond with the hydroxyl group of Y156 of TM3. Once stabilized, Y326 of TM7 sterically pushes TM6 outwards through interactions with L306 and L305 of TM6, resulting in coupled activation of G-protein binding (Figure 4D).

### G protein coupling and activation mechanism

PAR1 and PAR2 exhibit similar G-protein coupling profiles, both leading to strong Gαq-mediated signaling pathways. Structural alignments between Gαq in PAR1 and PAR2 complexes reveal notable conformational differences. The α5 helix of Gαq in both receptors shows a deflection of ~7° (Figure 5A-B), and a spatial separation of ~6.4 Å is observed in their respective αN helices (Figure 5C). These findings highlight the intricate and receptor-specific nature of GPCR-G protein interactions, emphasizing that insights from one receptor subtype cannot be directly transferred to another. Our study elucidates G-protein coupling specificities in PAR1 and PAR2, revealing the complex interplay of structural elements that govern these interactions.

**Figure 5.**
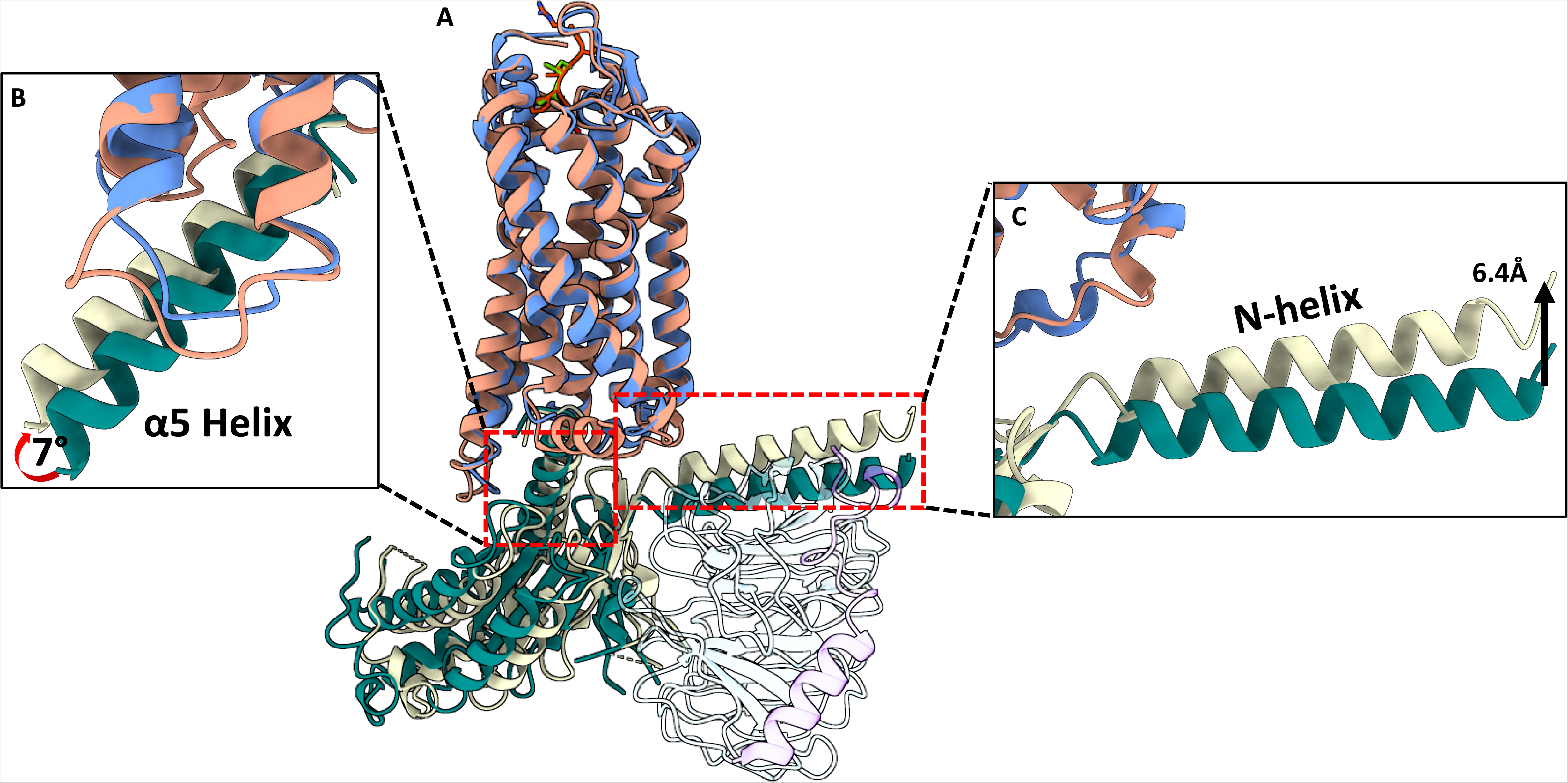
Interaction between PAR with Gq. **A**, Structural alignment between PAR1-Gαq and PAR2-Gαq. **B,C**, Comparative analysis of the Gαq conformation in the α5 helix (**B**) and αN domain (**C**).

Both PAR1 and PAR2 share a common activation mechanism involving structural changes upon ligand binding. In PAR1, the tethered ligand induces an outward movement of TM6 and shifts TM5, facilitating receptor activation by pushing down two critical tyrosines (Y350 and Y353), leading to TM6 movement.

Similarly, in PAR2, the tethered ligand also pushes down key tyrosines (Y323 and Y326), causing a conformational shift in TM6. These shifts are crucial for activating the G protein, highlighting the receptor-specific yet common activation dynamics across both PAR1 and PAR2 (Figure 6).

**Figure 6.**
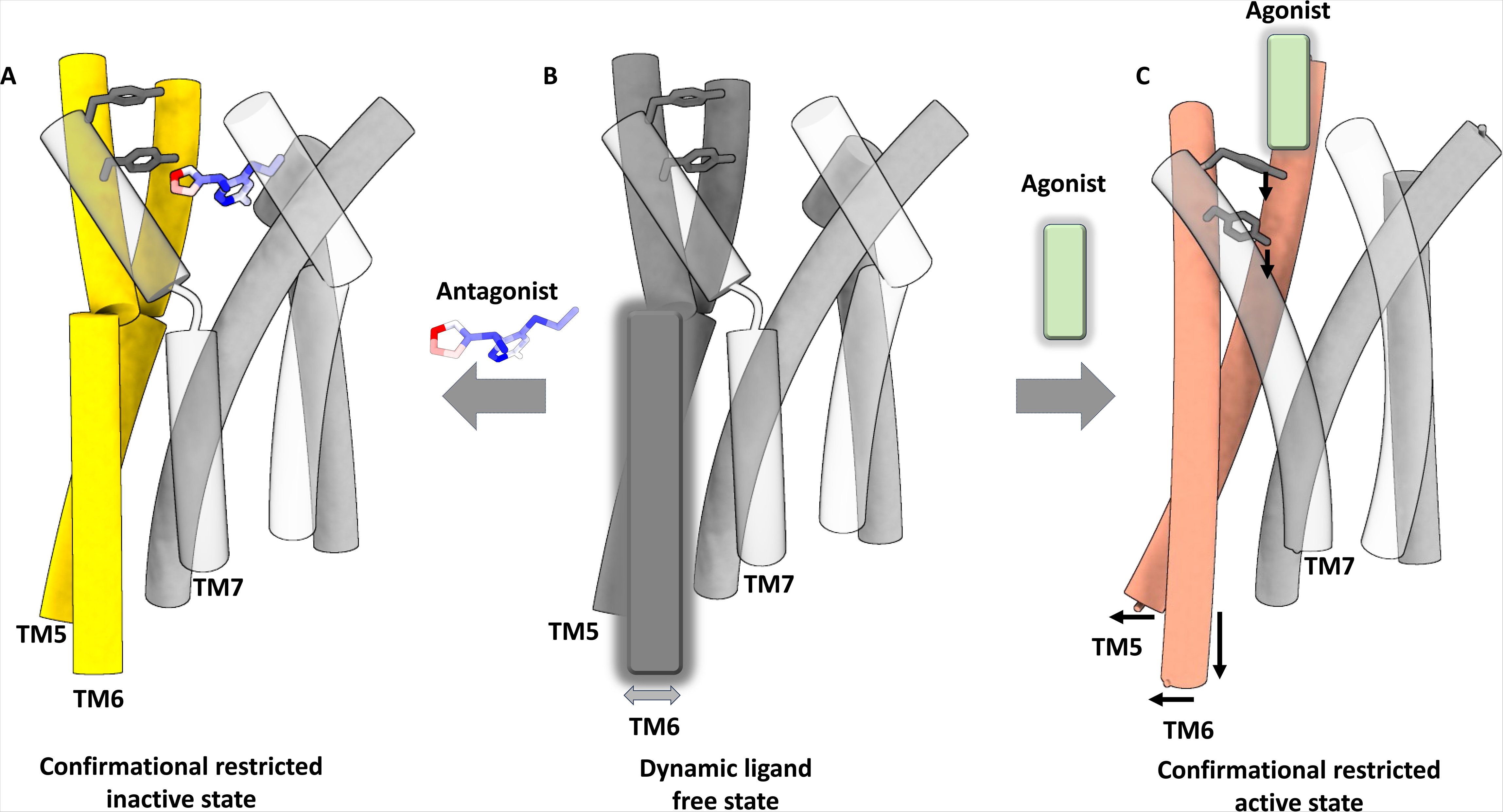
The activation mechanism of PAR1 and PAR2 activation and inactivation. Schematic representation of PAR1 and PAR2 in ligand-free (middle), antagonist-bound (left), and agonist-bound (right) states. The sidechain of two mechanistically important tyrosines are shown in stick representation. Conformational flexibility in TM7, TM5 and TM6 in the different state is indicated with black arrows, while conformational changes upon agonist and antagonist binding are highlighted.

## Discussion

In this study, we advanced our understanding of the structural mechanisms underlying the activation of PAR1 and PAR2 by their endogenous tethered ligand. Our cryo-EM structures provided a detailed view of the ligand-binding sites in PAR1 and PAR2, identifying the orthosteric binding pocket for the first time. Unlike previous findings[23], this binding pocket of these proteins are shallow and close to the extracellular side of the receptor. Due to its constricted nature, the binding of antagonists in PAR1 leads to significant conformational changes in TM5 and TM6 (Figure 3B).

The PAR2 inhibitor AZ8838, which binds much deeper than the tethered ligand, may function as a negative allosteric modulator (NAM). Unlike most Class A and Class B GPCRs, the movement of TM6 in PAR1 and PAR2 is relatively minor, indicating a unique G protein coupling mechanism. Both structures reveal a common activation mechanism, where the tethered ligand pushes down key tyrosines (Y323 and Y326, Figure 4D), both of which are conserved across all 4 human PARs, leading to TM6 movement. This structural information is crucial for developing true antagonists and designing PAR2-targeted therapeutics.

## Method

### Constructs

The human PAR1 and PAR2 gene were subcloned into the pFastBac plasmid with a prolactin-signal peptide sequence on its N-terminus and the LgBiT fused to its c-terminus followed by a Flag tag for purification. The first 41 residues (residue 1-41) of PAR1 and the first 36 residues (residue 1-36) were chopped off to mimic the protease cleavage. The HiBiT was fused to the c-terminus of human Gβ_1_ and cloned into pFastBac plasmid as described in the VIP1R paper. The N-terminus (residue 1-32) of human Gα_q_ was replaced by the N-terminus of G_i_ (residue 1–28) and subcloned into pFastBac plasmid. The wild-type human Gγ_2_ was cloned into pFastBac plasmid. The scFv16 that encodes the single-chain variable fragment of mAb16 was subcloned into pFastBac plasmid.

### Expression and purification of PAR1-Gαq and PAR2-Gαq complex

Bacmid preparation and virus production were performed according to the Bac-to-Bac baculovirus system manual (Gibco, Invitrogen). For expression, the *Spodoptera frugiperda* (Sf9) cells at density of 2 × 10^6^ cells per ml were co-infected with baculovirus encoding the PAR1-LgBiT-Flag, G_qiN,_ Gβ, Gγ, scFv16 and Ric8A protein at a ratio of 1:500 (virus volume vs cells volume). Cells were harvested 48 h after infection. Cell pellets were resuspended in 20 mM Hepes buffer (pH 7.5), 100 mM NaCl, and homogenized by douncing ~30 times. Apyase was added to the lysis at a final concentration of 0.5 mU/ml. To keep the complex stable, the lysate was incubated at room temperature for 1 h with flipping. Then, lauryl maltose neopentyl glycol (LMNG, Anatrace) was added at the final concentration of 0.5% to solubilize the membrane at 4 °C for 2 h. Then the lysis was ultracentrifuged at 56,000*g* (45,000 rpm) at 4 °C for 40 min. The supernatant was collected and incubated with anti-FLAG M2 resin for 2 h. The resin was washed with a buffer of 25 mM Hepes (pH 7.5), 100 mM NaCl and 0.001% LMNG, and 0.0005% cholesteryl hemi-succinate (CHS), then eluted with the same buffer plus 100 µb/ml FLAG peptide. The elution was concentrated and separated on a Superdex 200 Increase 10/300 GL (GE health science) gel infiltration column with a buffer of 25 mM Hepes (pH 7.5), 100 mM NaCl, and 0.001% LMNG. The peak corresponding to the PAR1-Gαq and PAR2-Gαq complex was concentrated at about 3 mg/ml and snap frozen for later cryo-EM grid preparation.

### Grid preparation and cryo-EM data collection

Three microliters PAR1-Gαq and PAR2-Gαq complex sample at ~3 mg/ml was applied to a glow-charged UltraFoil R1.2/1.3 grids (Quantifoil GmbH). The grids were vitrified in liquid ethane using Vitrobot Mark IV (Thermo Fisher Scientific) instrument in the setting of blot force of 5, blot time of 5 s, humidity of 100%, temperature of 4 °C. Single-particle cryo-EM data were collected on a Thermo Fisher Talos Arctica transmission electron microscope operating at 200 keV using parallel illumination conditions. Micrographs were collected using a Falcon 4i direct electron detector with a total electron exposure of 45 e^−^ Å^−2^ as EER format. All movie stacks were collected by the EPU program of FEI, nominal defocus value varied from 0.8 to 2.0 µm.

### Cryo-EM data processing

The cryoSPARC Live application of cryoSPARC v. 4 (Structura Biotechnology) was used to streamline the movie processing, CTF estimation, particle picking, and 2D classification. Preprocessing involved anisotropic motion correction and local CTF estimation. The data were curated by keeping only data with better than 4.5 Å determined by CTF fit resolution. Particle picking started with blob picking ~150 Å in diameter. Once a small set of particles was extracted and 2D class averages were obtained, the good 2D classes were used as templates for picking on the entire dataset. From the ~7 million (PAR1) and ~6 million (PAR2) particles initially picked, ~3.1 million and ~1.9 million particles were kept from the good 2D classes, respectively. Then ab initio 3D reconstruction was carried out in cryoSPARC v. 4 and one out of three classes was giving the reconstruction of the designed complex from ~1.9 million and ~0.4 million particles for PAR1 and PAR2, respectively. Homogenous refinement coupled with nonuniform and global CTF refinements gave the final 3D reconstructions at 2.74 Å (PAR1) and 3.1 Å (PAR2) resolution, respectively (0.143 gold standard FSC with correction of masking effects).

### Model building and refinement

Starting models for PAR1, PAR2, Gαq, Gβ_1_γ_2_, and scFv16 were based on Protein Data Bank (PDB) entries 3VW7[18], 5NDD[21], 6OIJ[24], respectively. All models were docked into the electron microscopy density map using UCSF Chimera[25]. The resulting model was subjected to iterative manual adjustment using Coot[26], followed by a Rosetta cryoEM refinement at relax model and Phenix real_space refinement[27]. The model statistics were validated using MolProbity. Structural figures were prepared in UCSF Chimera and PyMOL (https://pymol.org/2/). The statistics for data collection and refinement are included in Supplementary Table 1.

## Author Contributions

ZL, XL, and YK designed research. ZL, XL, and YK performed experiments. ZL, XL, GX, DC, VA, MF, NA, HZ, AM and YK analyzed data. ZL, XL, AM and YK wrote the manuscript.

---

**Supplemental Table 1.**

|  | **PAR1-Gαq** | **PAR2-Gαq** |
| --- | --- | --- |
| **Data collection and processing** |  |  |
| Magnification | 165K | 165K |
| Voltage (kV) | 200 | 200 |
| Electron exposure (e-/Å<sup>2</sup>) | 45 | 45 |
| Defocus range (μm) | -0.8 to -2.0 | -0.8 to -2.0 |
| Pixel size (Å) | 0.69 | 0.69 |
| Symmetry imposed | C1 | C1 |
| Initial particle images (no.) | 7,045,823 | 6,134,573 |
| Final particle images (no.) | 1,904,551 | 404,635 |
| Map resolution (Å) |  |  |
| FSC threshold | 0.143 | 0.143 |
| Map resolution (Å) | 2.74 | 3.1 |
| **Refinement** |  |  |
| Model resolution (Å) | 3.1 | 2.7 |
| FSC threshold | 0.5 | 0.5 |
| Model-Map CC (mask) | 0.73 | 0.73 |
| **Model composition** |  |  |
| Non-hydrogen atoms | 8378 | 8500 |
| Protein residues | 1075 | 1086 |
| B factors (Å<sup>2</sup>) |  |  |
| Protein | 64.62 | 76.05 |
| R.m.s. deviations |  |  |
| Bond lengths (Å) | 0.004 | 0.003 |
| Bond angles (Å) | 0.990 | 0.541 |
| **Validation** |  |  |
| MolProbity score | 1.8 | 2.06 |
| Clash score | 10.13 | 12.75 |
| Rotamer outliers (%) | 0.44 | 0.33 |
| Ramachandran plot |  |  |
| Favored (%) | 96.02 | 93.06 |
| Allowed (%) | 3.98 | 6.85 |

**Extended Data Figure 1.**
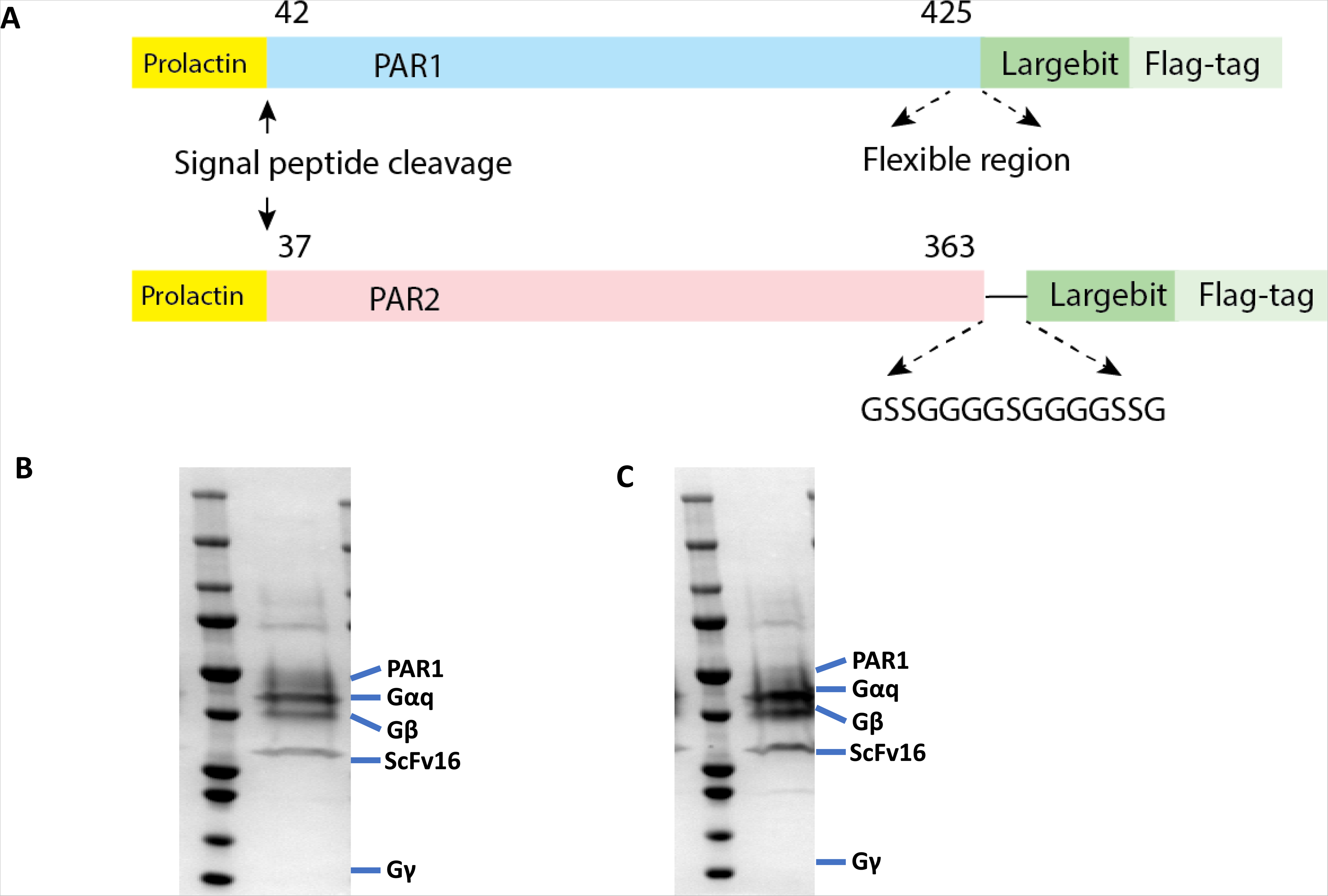
Construct design and protein purification. **A**, Cartoon representation of human PAR1 and PAR2 constructs. **B, C**, The final protein sample of PAR1-Gαq-scFv complex (**B**) and PAR1-Gαq-scFv complex (**C**).

**Extended Data Figure 2.**
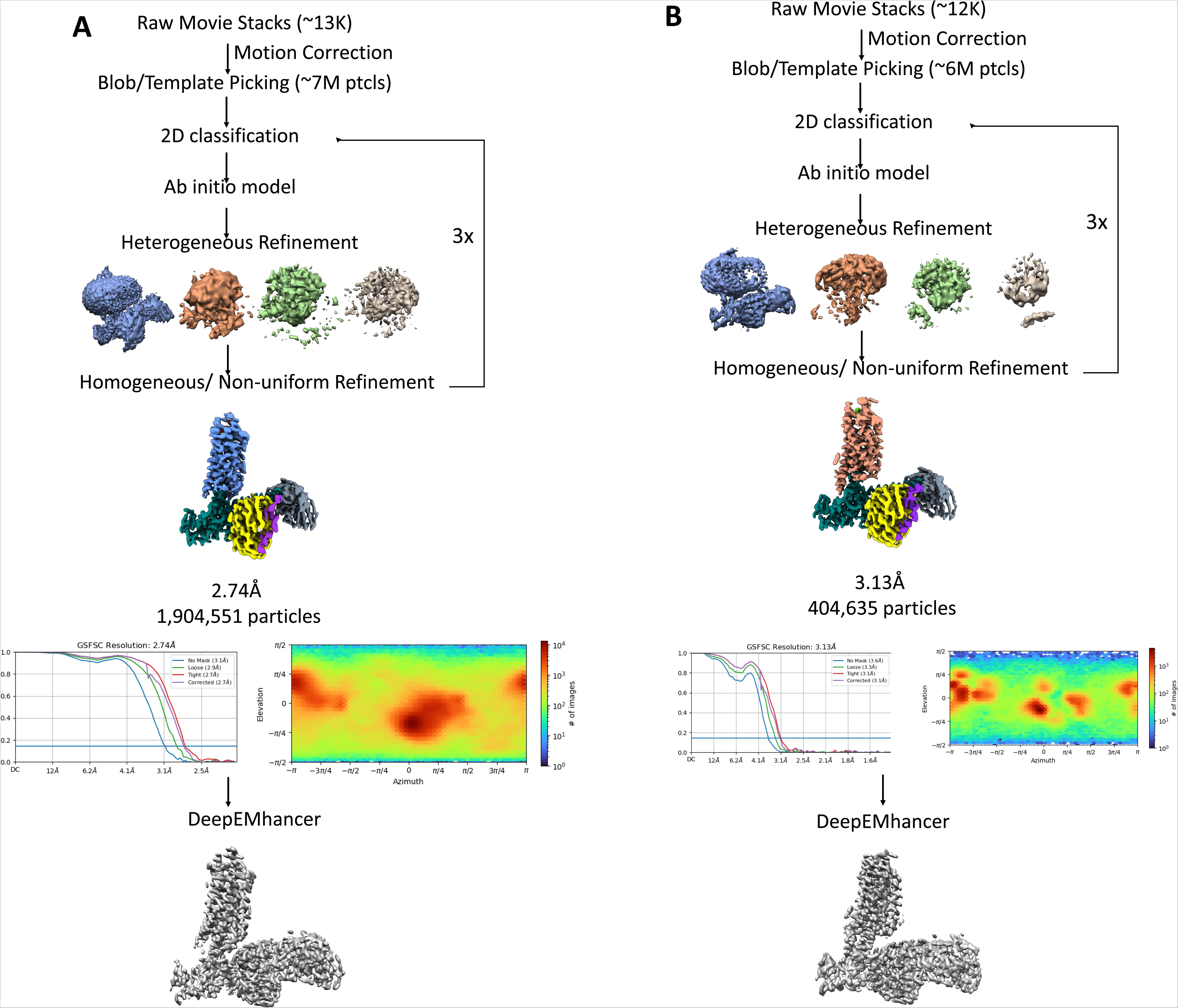
Cryo-EM analysis of PAR1-Gαq and PAR2-Gαq complexes. Image processing procedure of cryo-EM single particle analysis of the PAR1-Gαq (**A**) and PAR1-Gαq complex (**B**)

**Extended Data Figure 3.**
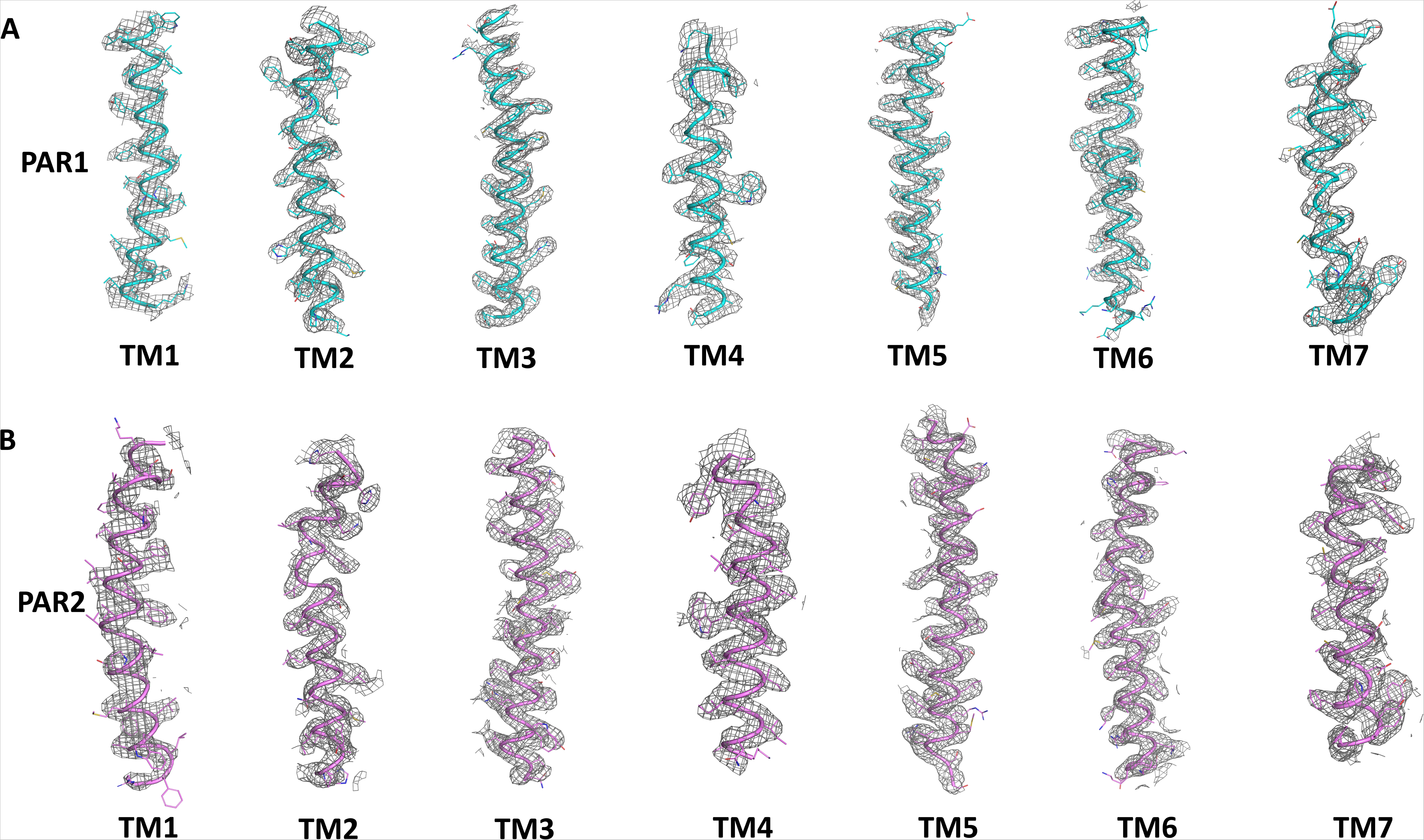
Density maps of PAR1-Gαq and PAR2-Gαq complexes. **A**, Depicts the representative resolution map of TM1-TM7 of PAR1 within PAR1-Gαq complex. **B**, Depicts the representative resolution map of TM1-TM7 of PAR1 within PAR1-Gαq complex.

**Extended Data Figure 4.**
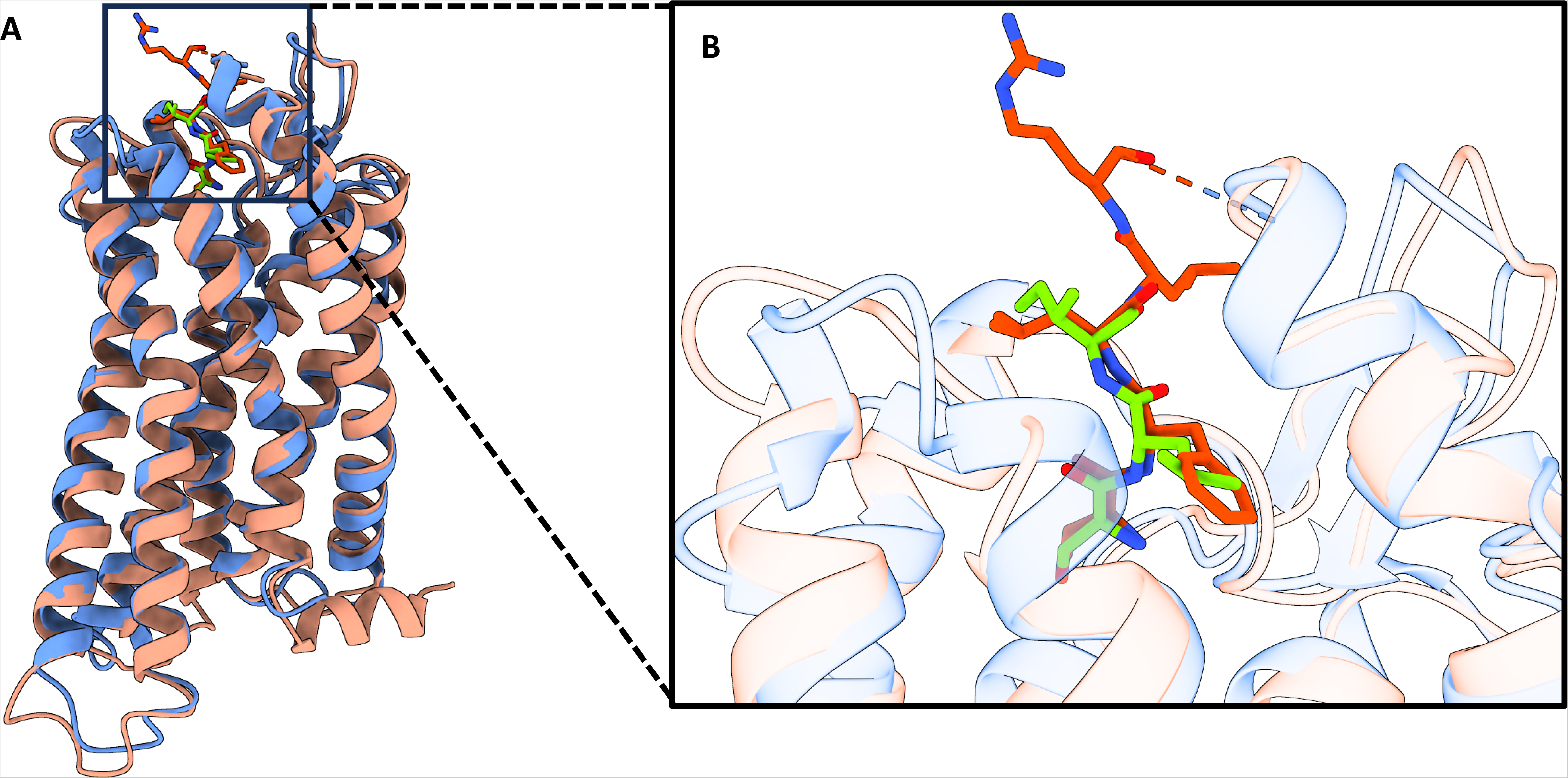
Confirmational comparison of PAR1 and PAR2. A, Structural superposition of the PAR1(Comflower) and PAR2 (Coral). B, Structural superposition of the tethered ligand of PAR1(Orange red) and the tethered peptide of PAR2 (Lime).

**Extended Data Figure 5.**
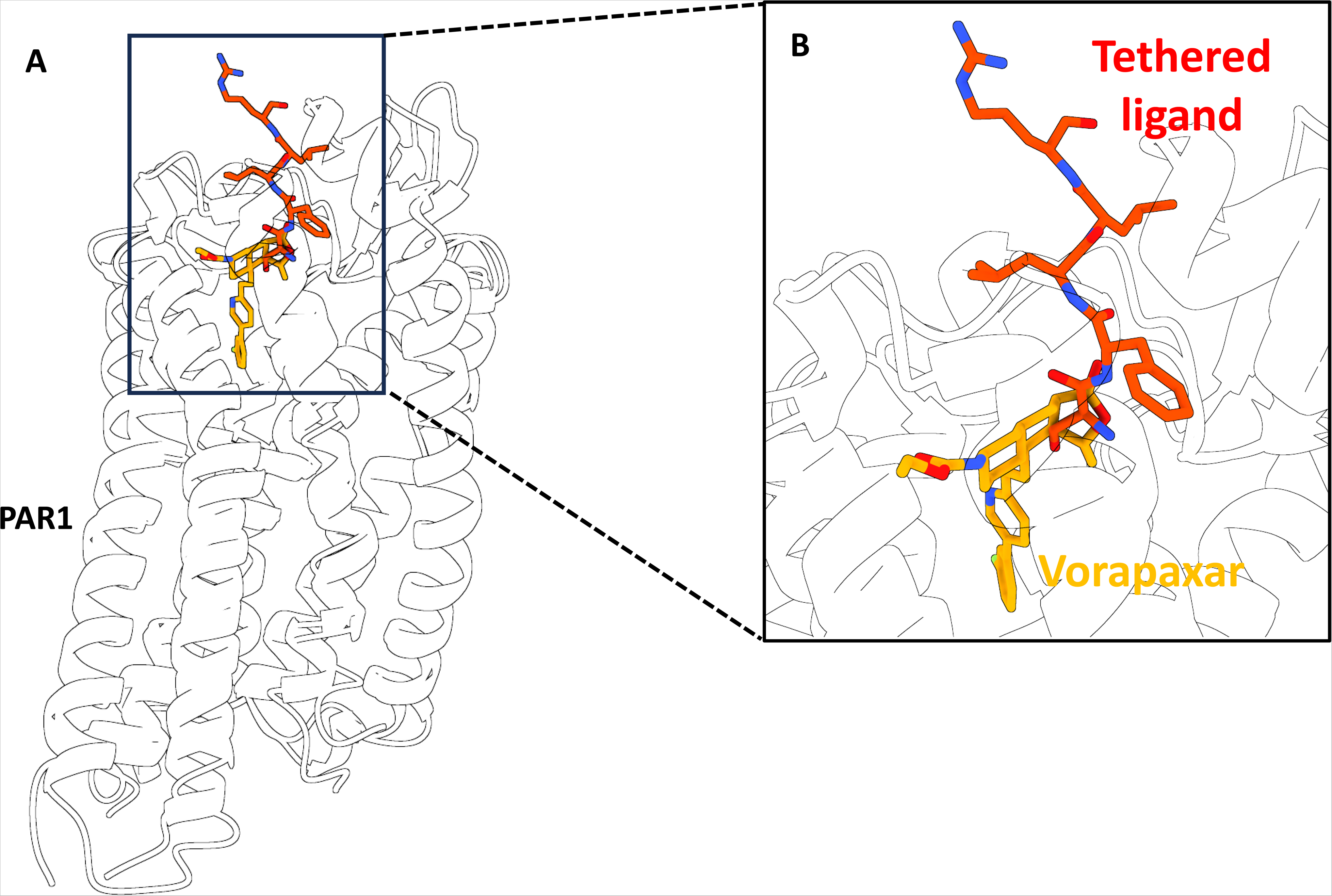
Structural comparison of active and inactive state of PAR1. Structural comparison shows that Vorapaxar (Orange) and the tethered peptide of PAR1 (Red) occupied the same binding site.

**Extended Data Figure 6.**
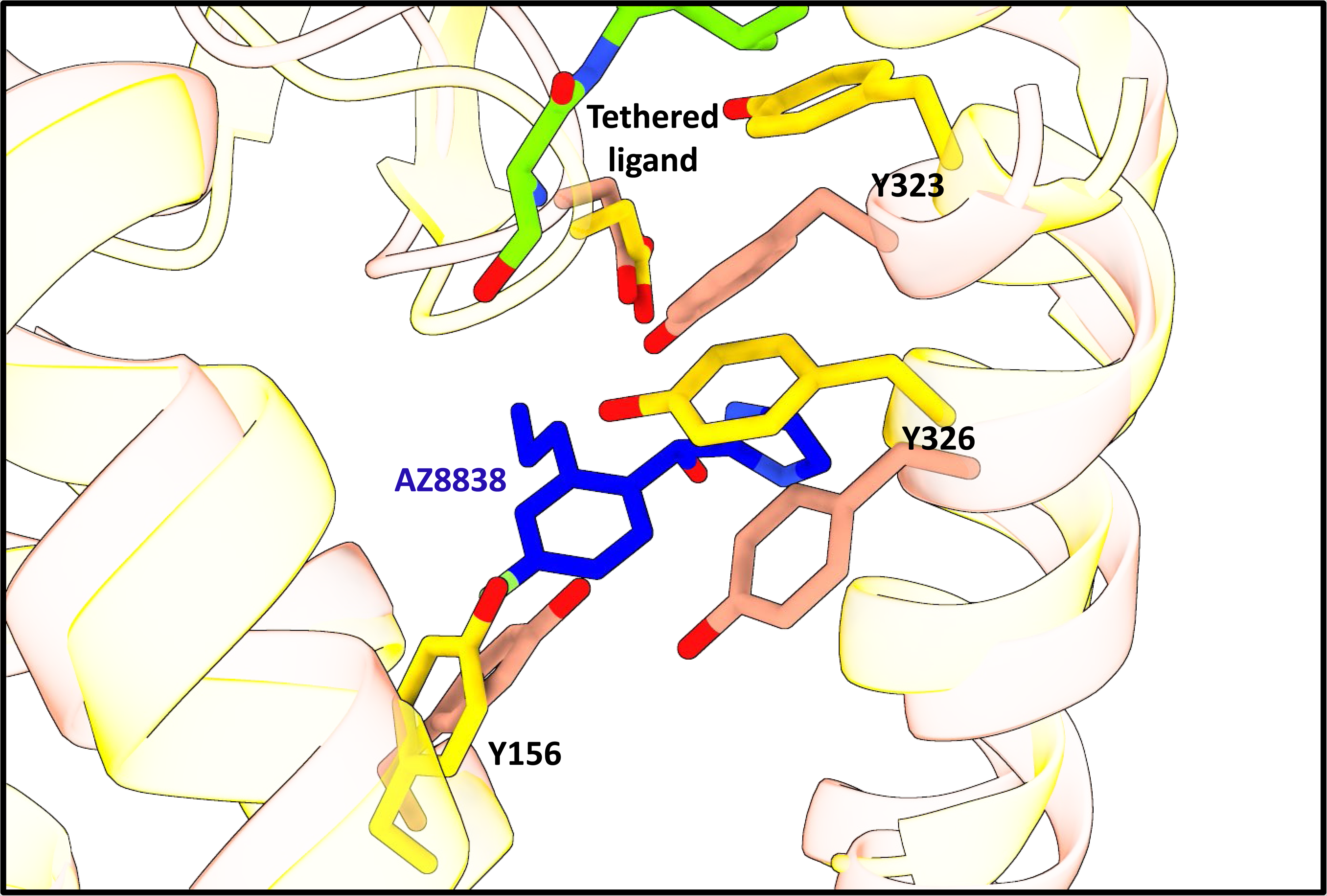
Confirmational comparison of active and inactive state of PAR2. The tethered ligand and AZ8838 occupy distinct binding pockets, resulting in significant conformational changes in several critical residues.

## References

1. Macfarlane, S.R., et al., Proteinase-activated receptors. Pharmacol Rev, 2001. 53(2): p. 245–82.

2. Helena Mangs, A. and B.J. Morris, The Human Pseudoautosomal Region (PAR): Origin, Function and Future. Curr Genomics, 2007. 8(2): p. 129–36.

3. Han, X., M.T. Nieman, and B.A. Kerlin, Protease-activated receptors: An illustrated review. Res Pract Thromb Haemost, 2021. 5(1): p. 17–26.

4. Han, X. and M.T. Nieman, The domino effect triggered by the tethered ligand of the protease activated receptors. Thromb Res, 2020. 196: p. 87–98.

5. Blackhart, B.D., et al., Ligand cross-reactivity within the protease-activated receptor family. J Biol Chem, 1996. 271(28): p. 16466–71.

6. Landis, R.C., Protease activated receptors: clinical relevance to hemostasis and inflammation. Hematol Oncol Clin North Am, 2007. 21(1): p. 103–13.

7. Grimsey, N.J. and J. Trejo, Integration of endothelial protease-activated receptor-1 inflammatory signaling by ubiquitin. Curr Opin Hematol, 2016. 23(3): p. 274–9.

8. Lin, H., et al., Cofactoring and dimerization of proteinase-activated receptors. Pharmacol Rev, 2013. 65(4): p. 1198–213.

9. Rattenholl, A. and M. Steinhoff, Proteinase-activated receptor-2 in the skin: receptor expression, activation and function during health and disease. Drug News Perspect, 2008. 21(7): p. 369–81.

10. Bar-Shavit, R., et al., Protease-activated receptors (PARs) in cancer: Novel biased signaling and targets for therapy. Methods Cell Biol, 2016. 132: p. 341–58.

11. Zhao, P., M. Metcalf, and N.W. Bunnett, Corrigendum: biased signaling of protease-activated receptors. Front Endocrinol (Lausanne), 2014. 5: p. 228.

12. Yau, M.K., L. Liu, and D.P. Fairlie, Toward drugs for protease-activated receptor 2 (PAR2). J Med Chem, 2013. 56(19): p. 7477–97.

13. Duan, J., et al., Cryo-EM structure of an activated VIP1 receptor-G protein complex revealed by a NanoBiT tethering strategy. Nat Commun, 2020. 11(1): p. 4121.

14. Xia, R., et al., Cryo-EM structure of the human histamine H(1) receptor/G(q) complex. Nat Commun, 2021. 12(1): p. 2086.

15. Ludeman, M.J., et al., PAR1 cleavage and signaling in response to activated protein C and thrombin. J Biol Chem, 2005. 280(13): p. 13122–8.

16. Liu, B., et al., The PAR2 signal peptide prevents premature receptor cleavage and activation. PLoS One, 2020. 15(2): p. e0222685.

17. Suen, J.Y., et al., Mapping transmembrane residues of proteinase activated receptor 2 (PAR(2)) that influence ligand-modulated calcium signaling. Pharmacol Res, 2017. 117: p. 328–342.

18. Zhang, C., et al., High-resolution crystal structure of human protease-activated receptor 1. Nature, 2012. 492(7429): p. 387–92.

19. Kang, Y., et al., Cryo-EM structure of human rhodopsin bound to an inhibitory G protein. Nature, 2018. 558(7711): p. 553–558.

20. Zhang, Y., et al., Cryo-EM structure of the activated GLP-1 receptor in complex with a G protein. Nature, 2017. 546(7657): p. 248–253.

21. Cheng, R.K.Y., et al., Structural insight into allosteric modulation of protease-activated receptor 2. Nature, 2017. 545(7652): p. 112–115.

22. Kennedy, A.J., et al., Protease-activated receptor-2 ligands reveal orthosteric and allosteric mechanisms of receptor inhibition. Commun Biol, 2020. 3(1): p. 782.

23. Kennedy, A.J., et al., Structural Characterization of Agonist Binding to Protease-Activated Receptor 2 through Mutagenesis and Computational Modeling. ACS Pharmacol Transl Sci, 2018. 1(2): p. 119–133.

24. Maeda, S., et al., Structures of the M1 and M2 muscarinic acetylcholine receptor/G-protein complexes. Science, 2019. 364(6440): p. 552–557.

25. Pettersen, E.F., et al., UCSF Chimera--a visualization system for exploratory research and analysis. J Comput Chem, 2004. 25(13): p. 1605–12.

26. Emsley, P. and K. Cowtan, Coot: model-building tools for molecular graphics. Acta Crystallogr D Biol Crystallogr, 2004. 60(Pt 12 Pt 1): p. 2126–32.

27. Adams, P.D., et al., PHENIX: a comprehensive Python-based system for macromolecular structure solution. Acta Crystallogr D Biol Crystallogr, 2010. 66(Pt 2): p. 213–21.

